# Whole-genome deep learning analysis reveals causal role of noncoding mutations in autism

**DOI:** 10.1101/319681

**Authors:** Jian Zhou, Christopher Y. Park, Chandra L. Theesfeld, Yuan Yuan, Kirsty Sawicka, Jennifer C. Darnell, Claudia Scheckel, John J Fak, Yoko Tajima, Robert B. Darnell, Olga G. Troyanskaya

## Abstract

We address the challenge of detecting the contribution of noncoding mutations to disease with a deep-learning-based framework that predicts specific regulatory effects and deleterious disease impact of genetic variants. Applying this framework to 1,790 Autism Spectrum Disorder (ASD) simplex families reveals autism disease causality of noncoding mutations by demonstrating that ASD probands harbor transcriptional (TRDs) and post-transcriptional (RRDs) regulation-disrupting mutations of significantly higher functional impact than unaffected siblings. Importantly, we detect this significant noncoding contribution at each level, transcriptional and post-transcriptional, independently and after multiple hypothesis correction. Further analysis suggests involvement of noncoding mutations in synaptic transmission and neuronal development, and reveals a convergent genetic landscape of coding and noncoding (TRD and RRD) *de novo* mutations in ASD. We demonstrate that sequences carrying prioritized proband *de novo* mutations possess transcriptional regulatory activity and drive expression differentially, and highlight a link between noncoding mutations and IQ heterogeneity in ASD probands. Our predictive genomics framework illuminates the role of noncoding mutations in ASD, prioritizes high impact transcriptional and post-transcriptional regulatory mutations for further study, and is broadly applicable to complex human diseases.

## Main

Great progress has been made in the past decade in discovering genetic causes of autism spectrum disorder, establishing *de novo* mutations, including copy number variants (CNVs) and point mutations that likely disrupt protein-coding genes, as an important cause of ASD^1,2^. Yet all known ASD-associated genes together explain a small fraction of new cases, and it is estimated that overall *de novo* protein coding mutations, including CNVs, contribute to only about 30% of simplex ASD cases^3^ (Supplementary Note 1). Despite the fact that the vast majority of the *de novo* mutations are located within intronic and intergenic regions, little is known with regard to the functions of these mutations and their contribution to the genetic architecture of disease in general, and ASD pathogenicity specifically.

A potential role of noncoding mutations in complex human diseases including ASD has long been speculated. Human regulatory regions show signs of negative selection^4^, suggesting mutations within these regions lead to deleterious effects, and studies of inherited common variants have shown enriched disease association in noncoding regions^5^. Furthermore, noncoding mutations affecting gene expression have been discovered to cause Mendelian diseases^6^ and shown to be enriched in cancer^7^. Expression dosage effects have also been suggested as underlying the link between CNVs and ASD^8^. Recently, parentally-inherited structural noncoding variants have been linked to ASD^9^. Also, on a small cohort of ASD families, some trends with limited sets of mutations have been reported^10–12^. Likewise, despite the major role RNA-binding proteins (RBPs) play in post-transcriptional regulation, little is known of the pathogenic effect of noncoding mutations affecting RBPs outside of the canonical splice sites. Thus, noncoding mutations could be a cause of ASD, yet no conclusive connection of regulatory *de novo* noncoding mutations, either transcriptional or post-transcriptional, to ASD etiology has been established.

Recent developments make it possible to perform large-scale studies that reliably identify noncoding *de novo* mutations at whole genome scale. The Simons Simplex Collection (SSC) whole genome sequencing (WGS) data for 1,790 families differs from many previous large-scale studies in design by including matched unaffected siblings^3,13–16^. These provide critical background controls for detecting excess of proband mutations, as it is otherwise hard to distinguish disease-relevant excess of mutations from irrelevant biological and technical variation, such as genetic background differences or artificial biases from sequencing, variant calling, and filtering procedures.

However, even with the study design with matched control individuals, detecting the *de novo* noncoding contribution is still challenging, and establishing the role of the vast noncoding space in the genetic basis of autism remains elusive. A recent study by Werling et al.^17^ in fact demonstrated that even when considering a wide variety of possible functional annotation categories (e.g. mutations in known regulatory sites, mutations at the location of known histone marks, mutations near ASD- or disease-relevant gene sets), no significant noncoding ASD-proband-specific signal was observed, and that approach would require a very large cohort to detect signal^17^. The challenge is to move beyond simple mutation counts, which are susceptible to both statistical power challenges and confounding factors, such as the rise in mutation counts with parental age^16,18^. This analytical challenge is shared in other psychiatric diseases with complex genetic bases, such as intellectual disabilities and schizophrenia. In fact, little is known about the contribution of noncoding rare variants or *de novo* mutations to human diseases beyond the less common cases with Mendelian inheritance patterns.

To address this challenge, we used a systematic approach (Fig. 1a) that reliably identifies impactful noncoding mutations, analogous to the genetic codon code which allows demarcation of protein coding mutations that likely disrupt protein function from synonymous ones. This enables comparison of mutational burden between probands and siblings not simply in terms of number of mutations, but in terms of their *functional impact*. Specifically, we used biochemical data demarcating DNA and RNA binding protein interactions to train and deploy a deep convolutional-neural-network-based framework that predicts the functional and disease impact of all 127,140 *de novo* noncoding mutations in the SSC, with independent models trained for DNA and RNA. Our framework estimates, with single nucleotide resolution, the quantitative impact of each variant on 2,002 specific transcriptional and 232 specific post-transcriptional regulatory features, including histone marks, transcription factors and RNA-binding protein (RBP) profiles.

**Fig. 1.**
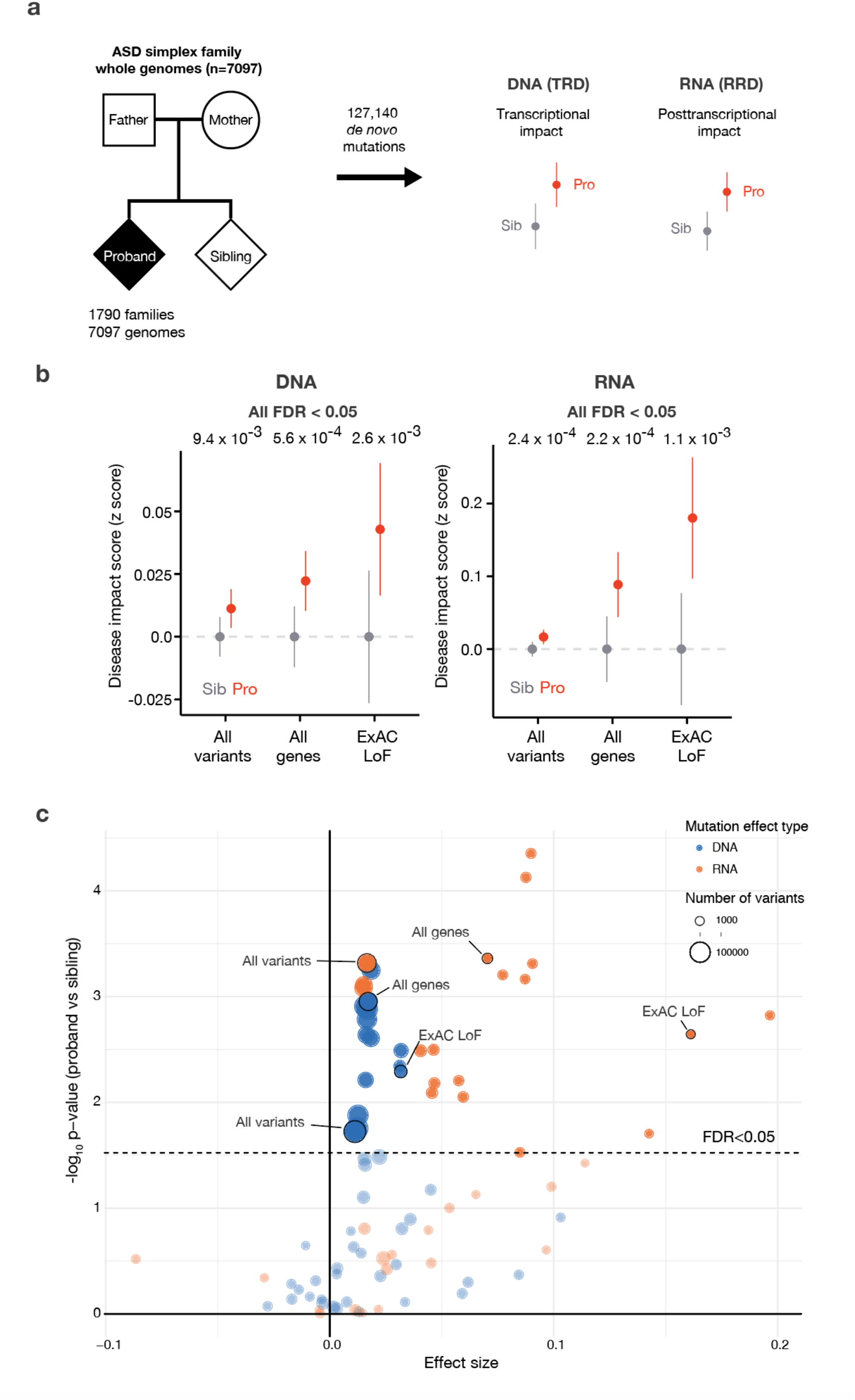
The elevated noncoding regulatory mutation burden in Autism Spectrum Disorder. a) Overall study design for deciphering the genome-wide *de novo* noncoding mutation contribution to ASD. 1,790 ASD simplex families whole genomes were sequenced to identify *de novo* mutations in the ASD probands and unaffected siblings. The identified 127,140 SNV *de novo* mutations were stratified by their predicted transcriptional (chromatin and TFs) and post-transcriptional (RNA-binding proteins) regulatory effect for comparison between probands and siblings. b) ASD probands possess mutations with significantly higher predicted disease impact scores compared to their unaffected siblings. We observe significant burden of both transcriptional (DNA - all variants, n=127,140) and post-transcriptional regulation (RNA - all transcribed variants, n=77,149) altering mutations in probands. This proband excess is stronger when restricted to mutation near all genes for DNA (n = 69,328) and near alternatively spliced exons for RNA (n =4,871), and even stronger near ExAC LoF intolerant (DNA n=14,873, RNA n=1,355) genes. For analyses that include gene sets, variants were associated with the closest gene within 100kb of the representative TSS for transcriptional regulatory disruption (TRD) analysis. For RNA regulatory disruption (RRD) analysis, variants located in the introns within 400bp of flanking exons in alternative splicing regulatory regions were used. Wilcoxon rank sum test (one-sided) was used for computing the significance levels. All predicted disease impact scores were normalized by subtracting average predicted disease impact scores of sibling mutations for each comparison (95% CI). Every result is significant with multiple hypothesis correction (FDR < 0.05). c) Genomic variant set analysis of noncoding burden for transcriptional- and posttranscriptional-disruptions. Significance level before and after correction for each category is listed in Supplementary Table 1. Categories in shown in Fig. 1b are annotated in figure. All gene lists were obtained from Werling et al.^17^. Distance cutoffs for DNA are 10kb, 50kb, 100kb, 500kb, ∞ to TSS, and distance cutoffs for RNA are 200bp, 400bp, ∞ to all exons or to all alternatively spliced exons. DNA results shown in orange and RNA in blue; dot size corresponds to number of variants in a category. Variant sets with >500 are displayed, full list results are available in Supplementary table 1. The dashed line indicates categories below FDR 0.05 threshold with the Benjamini-Hochberg method.

Using this approach, we discovered, independently at DNA and RNA regulation levels, a significantly (multiple-hypothesis corrected) elevated burden of disruptive transcriptional-regulatory disrupting (TRD) and RBP-regulatory disrupting (RRD) proband mutations in ASD, providing evidence for causality of noncoding regulatory *de novo* mutations in autism. Notably, the functional impact difference between proband and sibling mutations is significant when considering ***all** de novo* mutations, with elevated effect sizes observed around loss-of-function intolerant genes (ExAC^19^). We also identify specific pathways and tissues affected by these mutations, experimentally verify the differential regulatory effect of prioritized variants, and explore a link between the noncoding mutations and IQ in ASD.

## Results

### Contribution of transcriptional and post-transcriptional regulatory noncoding mutations to ASD

Analysis of noncoding effect contribution in ASD is challenging due to the difficulty of assessing which noncoding mutations are functional, and further, which of those contribute to the disease phenotype. For predicting the regulatory impact of noncoding mutations, we constructed a deep convolutional network-based framework to directly model the functional impact of each mutation and provide a biochemical interpretation including the disruption of transcription factor binding and chromatin mark establishment at the DNA level and of RBP binding at the RNA level (Supplementary Fig. 1). At the DNA level, the framework includes cell-type specific transcriptional regulatory effect models from over 2,000 genome-wide histone marks, transcription factor binding and chromatin accessibility profiles (from ENCODE and Roadmap Epigenomics projects^20,21^), extending the deep learning-based method that we described previously^10^ with redesigned architecture (leading to significantly improved performance, p=6.7×10^−123^, Wilcoxon rank-sum test). At the RNA level, our deep learning-based method was trained on the precise biochemical profiles of over 230 RBP-RNA interactions (CLIP) known to regulate a wide range of post-transcriptional regulation, including RNA splicing, localization and stability. At both transcriptional and post-transcriptional levels, our models are accurate and robust in whole chromosome holdout evaluations (Supplementary Fig. 1). Our models utilize a large sequence context to provide single nucleotide resolution to our predictions, while also capturing dependencies and interactions between various biochemical factors (e.g. histone marks or RBPs). This approach is data-driven, does not rely on known sequence information, such as transcription factor binding motifs, and it can predict impact of any mutation regardless of whether it has been previously observed, which is essential for the analysis of ASD *de novo* mutations. Finally, to link the biochemical disruption caused by a variant with phenotypic impact, we trained a regularized linear model using a set of curated human disease regulatory mutations^6^ (HGMD) and rare variants from healthy individuals in the 1000 Genomes populations^22^ to generate a predicted disease impact score for each autism mutation independently based on its predicted transcriptional and post-transcriptional regulatory effects.

With these approaches, we systematically assessed the functional impact of *de novo* mutations derived from 7,097 whole genomes from the SSC cohort with our framework (total 127,140 SNVs). When considering all *de novo* mutations, we observed a significantly higher functional impact in probands compared to unaffected siblings, independently at the transcriptional (p=9.4×10^−3^, one-side Wilcoxon rank-sum test for all; FDR=0.033, corrected for all mutation sets tested) and post-transcriptional (p=2.4×10^−4^, FDR=0.0049) levels (Fig. 1b, all variants). Recently, Werling et al.^17^ raised the challenge of detecting any significant proband-specific signal even with highly specific subsets of genes or genomic regions, and correspondingly emphasized the need for proper multiple hypothesis correction. Notably, our result above does not rely on any selection of variant subsets (e.g. those near predicted ASD-associated genes), is significant even after conservative multiple hypothesis correction, and, unlike the mutation counts, the predicted mutation effects are not correlated with parental age (Supplementary Fig. 2), a confounding factor of mutation count-based analysis.

To gain further insight into the ASD noncoding regulatory landscape, we conducted a comprehensive analysis, with full multiple hypothesis correction for all combinations of 14 gene-sets previously used in Werling et al.^17^ and 10 genomic regions tested (e.g. TSS or exon proximal). When restricted to genomic regions of higher regulatory potential (i.e. near TSS or alternatively spliced exons), we observed an increased dysregulation effect size (Fig. 1b-c, all genes, TRD p=5.6×10^−4^, FDR=0.0056; RRD p=2.2×10^−4^, FDR=0.0048). Among gene sets, we observed an elevated proband burden of high effect noncoding mutations close to loss-of-function (LoF) intolerant genes (pLI > 0.9 from ExAC, 3,230 genes, TRD p=2.6×10^−3^, FDR=0.013; RRD p=1.1×10^−3^, FDR=0.0078) (Fig. 1b-c, ExAC LoF), suggesting LoF intolerant genes are highly vulnerable to noncoding disruptive mutations in ASD. We observe these signals consistently across SSC cohort subsets that were sequenced in different phases (Supplementary Fig. 3). Importantly, we find the convergent signal across the noncoding genome in high regulatory and constrained gene regions independently in both RNA and DNA levels, providing further evidence of the casual role of noncoding variants in ASD (Full analysis p-values and FDRs are available in Supplementary Table 1).

### Tissue specificity and functional landscape of noncoding ASD-associated *de novo* mutations

Although one of the hallmarks of autism is altered brain development, a comprehensive tissue association has not been established for *de novo* noncoding variants. To explore the proband-specific tissue signal, we systematically tested the variant effects for tissue-specific genes derived from all 53 GTEx tissues and cell types^23^. We observed a consistent significant proband-specific mutation effect associated with brain tissues, with brain regions constituting the top 11 ranked tissues (by difference in proband vs sibling noncoding mutation effect) (Fig. 2a, all with FDR<0.05). This provides strong evidence that high impact variants from the noncoding genome of ASD probands likely disrupt brain-specific gene regulation, consistent with previous findings for protein coding mutations^28^.

**Fig. 2.**
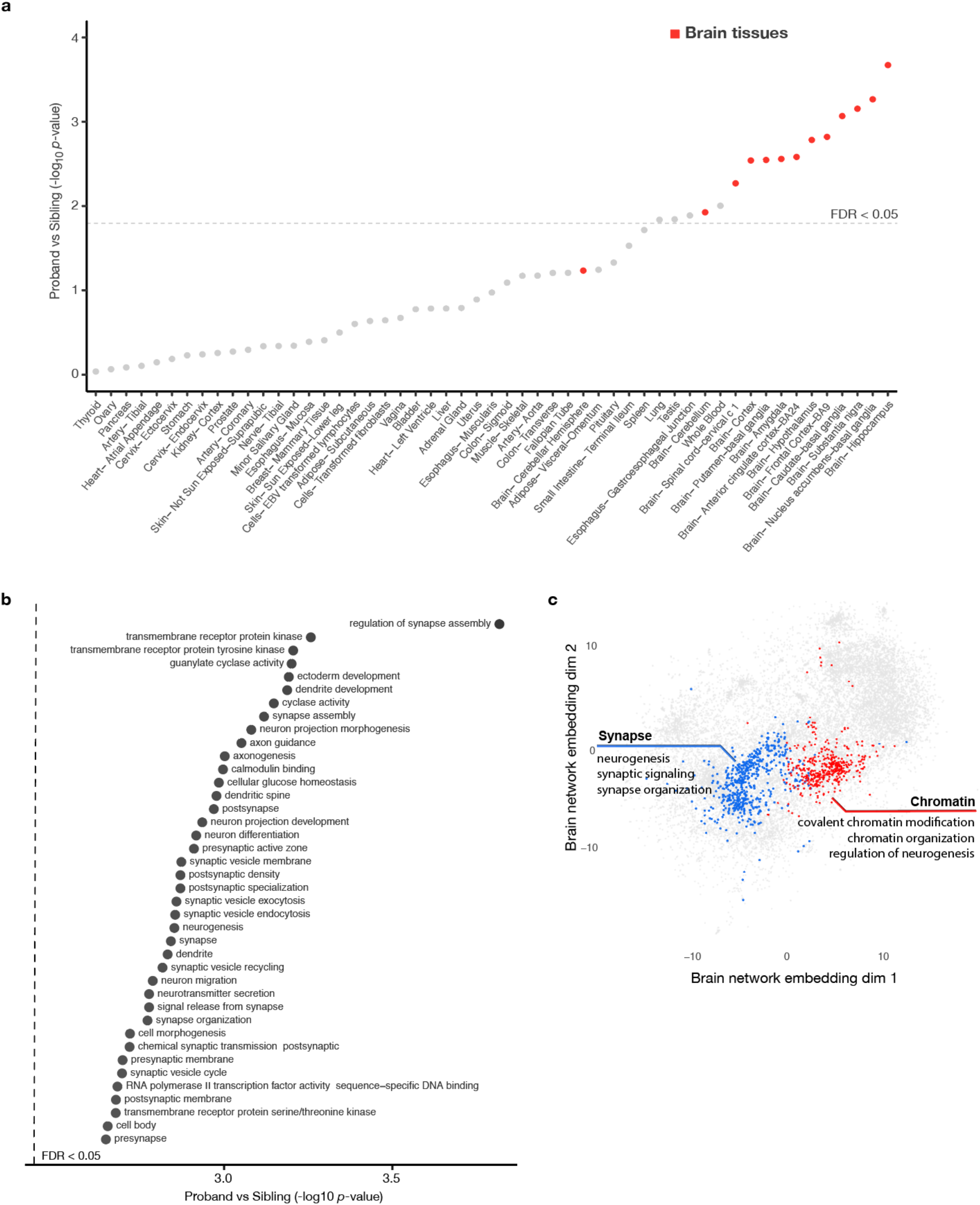
Analysis of noncoding mutations converges on brain specific signals and neurodevelopmental processes. a) Brain tissue-specific genes show strongest elevated proband-specific noncoding mutation burden. All 53 GTEx tissues are ranked by significance of increased proband mutation burden compared to unaffected siblings in tissue-specific genes (Methods). Dashed line indicates tissues below the FDR=0.05 threshold corrected with the Benjamini-Hochberg method. b) Neuronal function and development related processes show significant excess of proband mutation disease impact scores by NDEA (full list in Supplementary Table 3, see also Methods). The top processes (y-axis) and the p-values of proband excess (x-axis) are shown. All gene sets shown have FDR < 0.05. c) Genes with significant network neighborhood excess of high-impact proband mutations form two functionally coherent clusters (see annotations for representative enriched gene sets in each cluster, full list is in Supplementary Table 4). The brain functional network is visualized by computing two-dimensional embeddings with t-SNE (Methods). Genes, but not network edges, are shown for visualization clarity. The network differential enrichment analysis (NDEA) was performed on disease impact scores of all mutations within 100kb to representative TSSs (DNA) and all intronic mutations within 400nt to exon boundary (RNA). Clustering was performed with Louvain community clustering. All genes in the two clusters shown are with FDR < 0.1.

We next investigated the underlying processes and pathways impacted by *de novo* noncoding mutations in ASD. Such analysis is challenging because in addition to the variability in functional impact of mutations, ASD probands appear highly heterogeneous in underlying causal genetic perturbations^24^ and single mutations could cause a widespread effect on downstream genes. Thus to detect genes and pathways relevant to the pathogenicity of ASD TRD and RRD mutations, we developed a network-based statistical approach, NDEA (Network-neighborhood Differential Enrichment Analysis) (Supplementary Fig. 4). We used a brain-specific functional network that probabilistically integrates a large compendium of public omics data (e.g. expression, PPI, motifs) to represent how likely two genes are to act together in a biological process^25^. When applied to ASD *de novo* mutations, the NDEA approach identifies genes whose functional network neighborhood is significantly enriched for genes with stronger predicted disease impact in proband mutations compared to sibling mutations (Supplementary Table 2).

Globally, NDEA enrichment analysis pointed to a proband-specific role for noncoding mutations in affecting neuronal development, including in synaptic transmission and chromatin regulation (Fig. 2b), consistent with processes previously associated with ASD based on protein-coding variants^2^. Genes with significant NDEA enrichment were specifically involved in neurogenesis and grouped into two functionally coherent clusters with Louvain community detection algorithm (Fig. 2c). The synaptic cluster is enriched in ion channels and receptors involved in neurogenesis (p=5.6×10^−38^), synaptic signaling (p=4.8×10^−35^) and synapse organization (p=1.5×10^−18^), including previously known ASD-associated genes such as those involved in synapse organization SHANK2, NLGN2, NRXN2, synaptic signaling NTRK2 and NTRK3, ion channels CACNA1A/C/E/G, KCNQ2, and neurotransmission SYNGAP1, GABRB3, GRIA1, GRIN2A^26^. The synapse cluster is also significantly enriched for plasma membrane proteins (p=3.9×10^−24^). In contrast, the chromatin cluster, representing chromatin regulation related processes, displayed an overrepresentation of nucleoplasm (p=2.1×10^−9^) proteins, with diverse functional roles including covalent chromatin modification (p=2.5×10^−9^), chromatin organization(5.2×10^−8^) and regulation of neurogenesis (p=6.4×10^−5^). The chromatin cluster also includes many known ASD-associated genes such as chromatin remodeling protein CHD8, chromatin modifiers KMT2A, KDM6B, and Parkinson’s disease causal mutation gene PINK1^27^ which is also associated with ASD^26^. Overall, our results demonstrate pathway-level TRD and RRD mutation burden and identify distinct network level hot spots for high impact *de novo* mutations.

Next, we examined the genetic landscape of ASD-associated *de novo* noncoding and coding mutations. Specifically, in addition to the network analysis of noncoding mutations at the transcriptional and post-transitional level, we also applied it to the *de novo* coding mutations^2^. We compared the gene-specific NDEA statistic of elevated proband-specific noncoding mutation burden to that of the coding mutations, finding a significant positive correlation for both TRD and RRD (p=0.004 for TRD, p=0.042 for RRD; two-sided permutation test). Moreover, by network analysis, TRD and RRD are themselves significantly correlated (p=0.034 two-sided permutation test). This demonstrates a coding and noncoding mutations affect overlapping processes and pathways, indicating a convergent genetic landscape, and highlighting the potential of ASD gene discovery combining coding and noncoding mutations.

### Experimental study of ASD noncoding mutation effects on gene expression

Our analysis identified new candidate noncoding disease mutations with potential impact on ASD through regulation of gene expression. In order to add further evidence to a set of high confidence causal mutations, we experimentally studied allele-specific effects of predicted high-impact mutations in cell-based assays. Thirty four genomic regions showed strong transcriptional activity with 94% proband variants (32 variants) showing robust differential activity (Fig. 3, Methods); demonstrating that our prioritized *de novo* TRD mutations do indeed lie in regions with transcriptional regulatory potential and the predicted effects translate to measurable allele-specific expression effects. Among these genes with the demonstrated strong differential activity mutations, NEUROG1 is an important regulator of initiation of neuronal differentiation and in the NDEA analysis had significant network neighborhood proband excess (p=8.5×10^−4^), and DLGAP2 a guanylate kinase localized to the post-synaptic density in neurons. Mutations near HES1 and FEZF1 also carried significant differential effect on activator activities: neurogenin, HES, and FEZF family transcription factors act in concert during development, both receiving and sending inputs to Wnt and Notch signaling in the developing central nervous system and interestingly, the gut, to control stem cell fate decisions^28–32^; and Wnt and Notch pathways have been previously associated with autism^24,33^. SDC2 is a synaptic syndecan protein involved in dendritic spine formation and synaptic maturation, and a structural variant near the 3’ end of the gene was reported in an autistic individual (reviewed in Saied-Santiago, 2017^34^). Thus, our method identified alleles of high predicted impact that do indeed show changes in transcriptional regulatory activity in cells. Since many autism genes are under strong evolutionary selection, only effects exerted through (more subtle) gene expression changes may be observable because complete loss of function mutations may be lethal. This implies that further study of the prioritized noncoding regulatory mutations should yield insights into the range of dysregulations associated with autism.

**Fig. 3.**
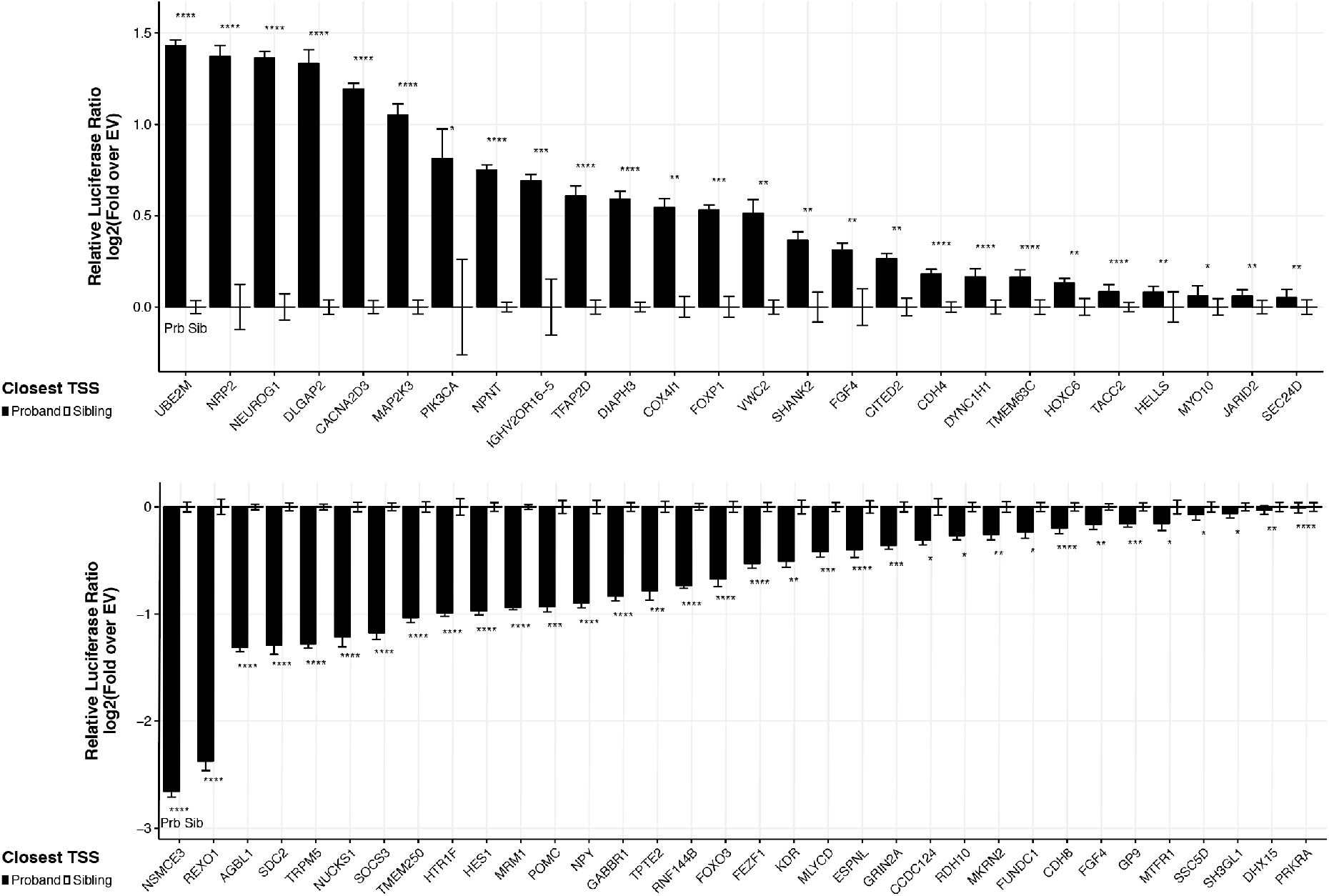
Allele-specific transcriptional activity of ASD noncoding mutations. Differential expression by proband or sibling alleles in a dual luciferase assay demonstrated that 32 predicted high disease impact mutations fall in active regulatory elements and the mutations confer substantial changes to the regulatory potential of the sequence. Y-axis shows the magnitude of transcription activation activity normalized to sibling allele. The error bar represents standard error of the mean. Significance levels were computed based on t-test (two-sided).

### Case study: association of IQ with *de novo* noncoding mutations in ASD individuals

*De novo* noncoding mutations provide a vast space for exploration of phenotype heterogeneity in ASD. To illustrate the potential of such analyses, we performed a case study focused on IQ. Intellectual disability is estimated to impact 40-60% of autistic children^35^, and ASD individuals can over-inherit common variants associated with high education attainment^36^. The genetic basis of this variation is not well understood. Despite the genetic complexity observed in association with ASD proband IQ, past efforts to identify mutations that contribute to ASD found that these mutations are also negatively correlated with IQ. Specifically, in analyses of exome sequencing data from different ASD cohorts, a significant association of higher burden of *de novo* coding likely-gene-disrupting (LGD) (also shown for WGS data in Supplementary Fig. 5) and large copy number variation (CNV) mutations with lower proband IQ was observed^2,8^. For *de novo* noncoding mutations analyzed in this study, we observe a significant association between noncoding mutations and IQ in ASD individuals. Intriguingly, we find that higher *IQ* ASD individuals have a higher burden of TRDs, whereas lower *IQ* ASD individuals have a higher burden of RRDs in ExAC LoF intolerant genes (Supplementary Fig. 6, DNA p=0.016, RNA p=0.020). Thus, it is tempting to speculate that while mutations that are damaging to the protein through disruption of coding (LGD or large CNVs) or RNA processing (RRD) are likely to increase the risk of lower IQ in ASD context, mutations affecting transcriptional regulation (TRDs) can affect ASD without the coupled negative effect on IQ.

### Conclusions

Even with great strides in understanding the causes of ASD by sequencing and phenotyping of multiple cohorts in the recent years, much of the genetic basis underlying autism remains undiscovered. While a number of coding variants have been associated with ASD, no systematic evidence of *de novo* noncoding effect has been observed. Here we present a novel deep-learning based approach for quantitatively assessing the impact of noncoding mutations on human disease. Our approach addresses the statistical challenge of detecting the contribution of noncoding mutations by predicting their specific effects on transcriptional and post-transcriptional levels. This approach is general and can be applied to study contributions of noncoding mutations to any complex disease or phenotype.

Here, we apply it to ASD using the 1,790 whole genome sequenced families from the Simons Simplex Collection, and for the first time demonstrate significant proband-specific signal in regulatory *de novo* noncoding space. Importantly, we independently detect this signal not only at the transcriptional level, but also find significant proband-specific RRD burden. Previously, there’s been limited evidence for disease contribution of mutations disrupting post-transcriptional mechanisms outside of the canonical splice sites. We demonstrate significant ASD disease association at the *de novo* mutation level for variants impacting a large collection of RBPs regulating post-transcriptional regulation. Overall, our results suggest that both transcriptional and posttranscriptional mechanisms play a significant role in ASD etiology and possibly other complex diseases.

Although previous work established that severe genetic perturbations such as CNVs and LGDs associated with ASD link to more severe intellectual disability and lower IQ, we show that this relationship may not be universal for all ASD causal genetic perturbations. This is important as it provides initial evidence that ASD and IQ can be genetically uncoupled at *de novo* mutation level.

Our analyses also demonstrate the potential of predicting disease phenotypes from genetic information including *de novo* noncoding mutations. We provide a resource for further research into understanding the mechanism of noncoding impact on ASD, including computationally prioritized TRD and RRD mutations with strong predicted regulatory effects, as well as potentially disease contributing ASD proband mutations with experimentally confirmed effects (Supplementary Table 5-6).

## Methods

### *De novo* mutation calling and filtering

The Simons Simplex Collection WGS data was made available via Simons Foundation Autism Research Initiative (SFARI), and was processed to generate variant calls via the standard GATK pipeline. To call *de novo* single nucleotide substitutions, inherited mutations were removed, and candidate *de novo* mutations were selected from the GATK variant calls where the alleles were not present in parents and the parents were homozygous with the same allele. DNMFilter classifier was then used to score each candidate *de novo* mutation and a threshold of probability > 0.75 was applied to phase 1 and 2 and a threshold of probability > 0.5 was applied to phase 3 to obtain a comparable number of high-confidence DNM calls across phases.

The DNMFilter^49^ classifier was trained with an expanded training set combining the original training standards with the verified DNMs from the pilot WGS studies for the 40 SSC families families^16^. *De novo* mutations calls within the repeat regions from RepeatMasker^50^ were removed. The WGS DNM calls were compared against exome sequencing *de novo* mutations calls and previously validated SSC *de novo* mutations^37^: 91.1% of the exome sequencing mutations calls and 93.2% of the validated mutations were rediscovered in our mutations calls. Further filtering was then applied to remove variants that were called in more than one SSC families.

### Training of DNA transcriptional regulatory effects and RNA posttranscriptional effects models

For training the transcriptional regulatory effects model, training labels, such as histone marks, transcription factors, and DNase I profiles, were processed from uniformly processed ENCODE and Roadmap Epigenomics data releases. The training procedure is as described in Zhou and Troyanskaya^21^ with the following modifications. The model architecture was extended to double the number of convolution layers for increased model depth (see Supplementary Note 2 for details). Input features were expanded to include all of the released Roadmap Epigenomics histone marks and DNase I profiles, resulting in 2,002 total features (Supplementary Table 7) compared to 919 original features.

For training the post-transcriptional regulatory effects model, we utilized the DeepSEA network architecture and training procedure with RNA-binding protein (RBP) profiles as training labels (full list of parameters used in model is in Supplementary Note 2). We uniformly process RNA features composed of 231 CLIP binding profiles for 82 unique RBPs (ENCODE and previously published CLIP datasets) and a branchpoint mapping profile as input features (full list of experimental features listed in Supplementary Table 8). CLIP data processing followed our previous detailed pipeline^38^, all CLIP peaks with p-value < 0.1 were used for training with an additional filter requirement of two-fold enrichment over input for ENCODE eCLIP data. In contrast to the DeepSEA, only transcribed genic regions were considered as training labels for the post-transcriptional regulatory effects model. Specifically, all gene regions defined by Ensembl (mouse build 80, human build 75) were split into 50nt bins in the transcribed strand sequence. For each sequence bin, RBP profiles that overlapped more than half were assigned a positive label for the corresponding RBP model. Negative labels for a given RBP model were assigned to sequence bins where other RBP’s non-overlapping peaks were observed. Note that our deep learning models, both transcriptional and post-transcriptional, does not use any mutation data for training, thus it can predict impacts for any mutation regardless of whether it has been previously observed.

### Disease impact score prediction

We used curated disease regulatory mutations and rare variants from healthy individuals to train a model that prioritizes likely disease-impacting mutations based on the predicted transcriptional or post-transcriptional regulatory impacts of these mutations. As positive examples, we used regulatory mutations curated in the Human Gene Mutation Database (HGMD). As negative examples of background mutations, we used rare variants that were only observed once within the healthy individuals from the 1000 Genomes project ^22^. Absolute predicted probability differences computed by the convolutional network transcriptional regulatory effects model (described above) were used as input features for each of the 2,002 transcriptional regulatory features and for the 232 post-transcriptional regulatory features in the disease impact model. Input features were standardized to unit variance and zero mean before being used for training. We separately trained a L2 regularized logistic regression model for transcriptional effect model (lambda=10) and post-transcriptional effect model (lambda=10, using only genic region variant examples) with the xgboost package (https://github.com/dmlc/xgboost).

### Gene sets and resources

All gene sets used are from Werling et al.^17^. The 14 gene-sets include GENCODE protein coding genes, Antisense, lincRNAs, Pseudogenes, genes with loss-of-function intolerance (pLI) score > 0.9 from ExAC^19^, predicted ASD risk genes (FDR < 0.3) from Sanders et al.^8^, FMRP target genes^39^, Genes associated with developmental delay^40,41^ and CHD8 target genes^42,43^. For genes with expression specific to each 53 GTEx tissue, we used expression table from GTEx 1.8 (gene median TPM per tissue)^23^, we selected genes for which expression in a given tissue was five times higher than the median expression across all tissues.

We determined the representative TSS for each gene based on FANTOM CAGE transcription initiation counts relative to GENCODE gene models. Specifically, a CAGE peak is associated to a GENCODE gene if it is within 1000bp from a GENCODE v24 annotated transcription start site^44,45^. Peaks within 1000bp to rRNA, snRNA, snoRNA or tRNA genes were removed to avoid confusion. Next, we selected the most abundant CAGE peak for each gene, and took the TSS position reported for the CAGE peak as the selected representative TSS for the Gene. For genes with no CAGE peaks assigned, we kept the GENCODE annotated gene start position as the representative TSS. FANTOM CAGE peak abundance data were downloaded at http://fantom.gsc.riken.jp/5/datafiles/latest/extra/CAGE_peaks/ and the CAGE read counts were aggregated over all FANTOM 5 tissue or cell types. GENCODE v24 annotation lifted to GRCh37 coordinates were downloaded from http://www.gencodegenes.org/releases/24lift37.html. All chromatin profiles used from ENCODE and Roadmap Epigenomics projects were listed in **Supplementary Table 7**. The HGMD mutations are from HGMD professional version 2018.1.

Human exons that are alternatively spliced (AS) were obtained from a recent study that has examined publicly available human RNA-seq data to annotate an extensive catalog of AS events^46^. Internal exon regions (both 5’SS & 3’SS flanking introns), upstream exon (5’SS flanking introns), and downstream terminal exon (3’SS flanking introns) were used for alternative exon definition types of cassette, mutually exclusive, tandem cassette exons. Terminal exon region was used for intron retention, alternative 3’ or 5’ exon AS exon types. All selected exon-flanking intronic regions were collapsed into a final set of genomic intervals used to subset SNVs that are located within alternative splicing exon region (200 or 400nts from exon boundary), illustrated in **Supplementary Fig. 7**.

### Network differential enrichment analysis (NDEA)

Brain-specific functional relationship networks integrate a wide-range of functional genomic data in a tissue-specific manner and predicted the probability of functional association between any pair of genes^25^. This network was filtered to only include edges with >0.01 probability (above Bayesian prior) to reduce the impact of noisy low-confidence edges.

For each gene *i*, we designed the neighborhood excess significance test which is a specific form of weighted t-test, specifically the t statistic is computed by

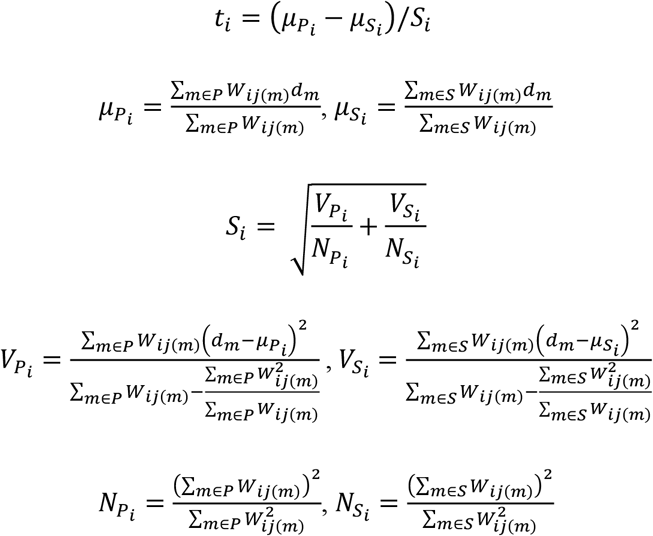

in which *μ_P_i__* and *μ_S_i__* are weighted averages of disease impact scores *d_m_* of all proband mutations *P* or all sibling mutations *S. W*_*ij*(*m*)_ is the network edge score (interpreted as functional relationship probability) between gene *i* and gene *j*(*m*) divided by the number of proband (if *m* is a proband mutation) or sibling (if *m* is a sibling mutation) mutations gene *j*(*m*) is associated to, where *j*(*m*) indicate the implicated gene of the mutation *m. P* and *S* are the set of all proband mutations and the set of all sibling mutations included in the analysis. *V_P_i__* and *V_S_i__* are the unbiased estimates of population variance of *μ_P_i__* and *μ_S_i__. N_P_i__* and *N_S_i__* are the effective sample sizes of proband and sibling mutations after network-based weighting for gene i.

Under null hypothesis of the two groups have no difference, the above t statistic approximately follows a t-distribution with the following degree of freedom:

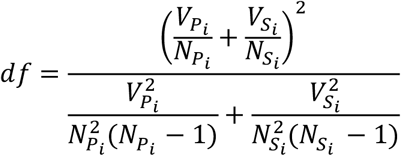

For testing significance difference between proband and sibling mutations, mutations within 100kb of the representative TSS of all genes and all intronic mutations within 400bp to exon boundary were included in this analysis. RNA model disease impact score z-scores were used as the mutation score for intronic mutations within 400bp to exon boundary and DNA model disease impact score z-scores were used for other mutations.

For gene set level NDEA, we create a meta-node that represents all genes that are annotated to the gene set (e.g. GO term). Then, the average of network edge scores for all genes in the meta-node is used as the weights to any given gene not part of the gene set. GO term annotations were pooled from human (EBI 5/9/2017), mouse (MGI 5/26/2017) and rat (RGD 4/8/2017). Query GO terms were obtained from the merged set of curated GO consortium^47^ slims from Generic, Synapse, ChEMBL, and supplemented by PANTHER^48^ GO-slim and terms from NIGO^49^.

For network-based analysis of correlation between coding and noncoding TRD and RRD mutations, we first compute the NDEA t-statistic for every gene for all protein *coding* mutations from SSC exome sequencing study^2,8^, all SSC WGS noncoding mutations within 100kb to a gene, and all SSC WGS genic noncoding mutations within 400bp to an exon, respectively. We then compute correlation across all resulting gene-specific t-statistics between all three pairs of mutation types. For testing statistical significance of the correlation, we permuted proband and sibling labels for all mutations to compute the null distributions of correlations for each pair of mutation type. 1000 permutations were performed.

### Network visualization and clustering

For network visualization, we computed a two-dimensional embedding with t-SNE^50^ by directly taking a distance matrix of all pairs of genes as the input. The distance matrix was computed as −log(probability) from the edge probability score matrix in the brain-specific functional relationship network. The Barnes-Hut t-SNE algorithm implemented in the Rtsne package was used for the computation. Louvain community clustering were performed on the subnetwork containing all protein-coding genes with top 10% NDEA FDR.

### Cloning of Variant Allele Genomic Regions

All genomic sequences were retrieved from the hg19 human genome assembly. For experimental testing, we selected variants of high predicted disease impact scores larger than 0.5 and included mutations near genes with evidence for ASD association, including those with LGD mutations (e.g. CACNA2D3) and a proximal structural variant (e.g. SDC2). For each allele (sibling or proband), we either cloned 230 nucleotides of genomic sequence amplified from proband lymphoblastoid cell lines or used fragments synthesized by Genewiz (**Supplementary Table 6**). In both cases, 15 nucleotide flanks on 5’ and 3’ ends matched each flank of the plasmid cloning sites. The 5’ sequence was TGGCCGGTACCTGAG and the 3’ sequence was ATCAAGATCTGGCCT. Synthesized fragments were cut with KpnI and BglII and cloned into pGL4.23 (Promega) cut with the same enzymes. PCR-amplified genomic DNA was cloned into pGL4.23 blunt-end cut with EcoRV and Eco53kI using GeneArtCloning method from Thermofisher Scientific. All constructs were verified by Sanger sequencing.

### Luciferase Reporter Assays

Human neuroblastoma BE(2)-C cells were plated at 2×10^4^ cells/well in 96-well plates and 24 hours later were transfected with Lipofectamine 3000 (L3000-015, Thermofisher Scientific) together with 75ng of Promega pGL4.23 firefly luciferase vector containing the 230nt of human genomic DNA from the loci of interest (**Supplementary Table 6**), and 4ng of pNL3.1 NanoLuc (shrimp luciferase) plasmid, for normalization of transfection conditions. 42 hours after transfection, luminescence was detected with the Promega NanoGlo Dual Luciferase assay system (N1630) and BioTek Synergy plate reader. Four to six replicates per variant were tested in each experiment. For each sequence tested, the ratio of firefly luminescence (ASD allele) to NanoLuc luminescence (transfection control) was calculated and then normalized to empty vector (pGL4.23 with no insert). Statistics were calculated from fold over empty vector values from each biological replicate. High-confidence differentially-expressing alleles were defined by their ability to show the same effect in each biological replicate (n=3, minimum), drive higher than control empty-vector level gene expression, and the two alleles had significantly different level of luciferase activity by two-sided t-test. For presentation of the data, we normalized the fold over empty vector value of the proband allele to that of the sibling allele.

### Transcriptional and post-transcriptional effect association with IQ

To analyze the association between transcriptional or post-transcriptional effect with IQ, we computed the maximum probability differences across features for each mutation, and tested for its association with IQ using linear regression with two-sided Wald test on the slope coefficient. For DNA analysis, we use all variants that are within 100kb from the TSS. For RNA analysis, we restrict the mutations to genes with ExAC pLI >0.9 and are intronic within 400nts to an exon in an alternatively splicing regulatory region.

